# DMA-tudor interaction modules control the specificity of *in vivo* condensates

**DOI:** 10.1101/2020.09.15.297994

**Authors:** Edward M. Courchaine, Andrew E.S. Barentine, Korinna Straube, Joerg Bewersdorf, Karla M. Neugebauer

## Abstract

Biomolecular condensation is a widespread mechanism of cellular compartmentalization. Because the ‘survival of motor neuron protein’ (SMN) is required for the formation of three different membraneless organelles (MLOs), we hypothesized that at least one region of SMN employs a unifying mechanism of condensation. Unexpectedly, we show here that SMN’s globular tudor domain was sufficient for dimerization-induced condensation *in vivo*, while its two intrinsically disordered regions (IDRs) were not. The condensate-forming property of the SMN tudor domain required binding to its ligand, dimethylarginine (DMA), and was shared by at least seven additional tudor domains in six different proteins. Remarkably, asymmetric versus symmetric DMA determined whether two distinct nuclear MLOs – gems and Cajal bodies – were separate or overlapping. These findings show that the combination of a tudor domain bound to its DMA ligand – DMA-tudor – represents a versatile yet specific interaction module that regulates MLO assembly and defines their composition.

## Introduction

In the past ten years, molecular condensation of proteins and RNA has emerged as a prominent mode of subcellular organization (Corbet and Parker, 2020; Courchaine et al., 2016; Shin and Brangwynne, 2017). Our understanding of macromolecular structures formed by condensation has developed from descriptive characterization of organelles like P-granules and nucleoli to mechanistic models of how specific molecules promote phase transitions (Brangwynne et al., 2009; Brangwynne et al., 2011; Pak et al., 2016; Wang et al., 2018). While the theoretical model of condensation has proven useful in many cases (A and Weber, 2019). The field lacks a complete understanding of how that conceptual framework accounts for the specificity, selectivity, and function of membraneless organelles (MLOs) (A and Weber, 2019; Alberti et al., 2019; Banani et al., 2016; McSwiggen et al., 2019). For example, several MLOs in the cell nucleus – nucleoli, Cajal bodies, gems, and histone locus bodies –show considerable overlap in composition but nevertheless do not fuse into one body, instead maintaining their independence (Machyna et al., 2013; Strom and Brangwynne, 2019).

Several molecular properties are now known to promote the formation of condensates. Early *in vitro* work established that multivalent binding is a key attribute of condensate formation, and those results have led to a model of Arp2/3 actin regulation condensates (Case et al., 2019; Li et al., 2012). Intrinsically disordered regions (IDRs) were also implicated in condensate formation and have since been the primary focus of many studies (Kato et al., 2012; Kwon et al., 2013). The capacity for individual amino acid residues of IDRs to act as multivalent interaction sites may provide a general explanation for why they readily phase separate *in vitro* (Lin et al., 2015; Nott et al., 2015; Patel et al., 2015; Sheu-Gruttadauria and MacRae, 2018; Wang et al., 2018). However, these studies have not given way to a universally applicable model of IDR-driven condensation in living cells. Indeed, the IDR of G3BP1 was recently reported to antagonize condensation of stress granules (Guillen-Boixet et al., 2020). The volume of *in vitro* data and sometimes conflicting models for biomolecular condensation underscores the need to address the mechanisms governing the specificity of condensate formation in the context of *in vivo* endogenous MLOs.

This study focuses on the ‘survival of motor neuron protein’ (SMN), which is an essential component in the biogenesis of small nuclear ribonucleoproteins (snRNPs) that are required for splicing (Pellizzoni et al., 1998; Zhang et al., 2011). SMN deficiency results in the fatal childhood disease, spinal muscular atrophy (SMA); yet SMN’s snRNP biogenesis role does not adequately explain the disease phenotype, because splicing is a required activity of every living cell (Buhler et al., 1999; Pellizzoni et al., 1998). SMN depletion results in the loss of three membraneless compartments – gems, Cajal bodies, and U-bodies (Girard et al., 2006; Lee et al., 2009; Lemm et al., 2006; Raimer et al., 2017; Shpargel et al., 2003; Strzelecka et al., 2010a). Cytoplasmic SMN promotes the assembly of the Sm ring on snRNPs but its molecular role after import to the nucleus is less clear (Buhler et al., 1999; Meister et al., 2001; Renvoise et al., 2006). How loss of cellular compartmentalization may contribute to SMA is currently unknown, but compromised Cajal body integrity has been reported in patient tissue (Tapia et al., 2012). We therefore sought to understand how SMN supports the assembly of diverse MLOs.

Here, we examine the role of SMN in the formation of MLOs by isolating each region of the SMN protein to assess its condensation potential. To do so, we repurposed the ‘optodroplet’ assay in which light is used to induce dimerization of protein domain candidates for condensation (Shin et al., 2017). Unexpectedly, we found that the critical domain for condensation of SMN was its globular tudor domain, not its IDRs. We developed a new analysis of optodroplet data in order to make quantitative comparisons between condensation that accounts for the expression level of each protein. This allowed us to show that dimerization induced condensation of the SMN tudor domain depends on binding to the post-translational modification dimethyl arginine (DMA), and that this property is shared amongst numerous additional tudor domains. Finally, using specific inhibitors of DMA synthesis, we found that it modulates the specific composition of two endogenous MLOs, the Cajal body and the gem. Our structure-function analysis of MLO formation reveals that the DMA-tudor module defines the specific composition of certain *in vivo* condensate MLOs.

## Results

SMN has three regions as distinguished by overall secondary structure (Figure 1A). N-terminal and C-terminal IDRs flank a single tudor domain that binds symmetric DMA on snRNP proteins and other ligands (Friesen et al., 2001; Wang and Dreyfuss, 2001; Zhang et al., 2011). The N-terminus contains a lysine-rich region, while the C-terminus has a proline-rich tract and a tyrosine/glycine repeat motif. Past work has shown that IDRs enriched for polar and aromatic amino acids can promote condensation, making SMN’s IDRs good candidates (Kato et al., 2012; Kwon et al., 2013; Wang et al., 2018). While some IDRs can form droplets spontaneously *in vitro*, they often require multimerization to overcome nonspecific interactions and condense *in vivo* (Protter et al., 2018; Shin et al., 2017). Thus, we fused each of the three SMN regions to the light- activated dimerization domain Cry2. This approach was previously exploited to form light-dependent ‘optodroplets’ with IDRs expressed in NIH-3T3 cells (Figure 1B) (Shin et al., 2017). In agreement with prior data, mCherry-Cry2 did not form clusters, whereas FUS^IDR^ and hnRNP- A1^IDR^ did form clusters upon light-induced dimerization (Figure S1A). We used this system to determine whether any portion of SMN directly promotes condensate formation *in vivo*.

**Figure 1.**
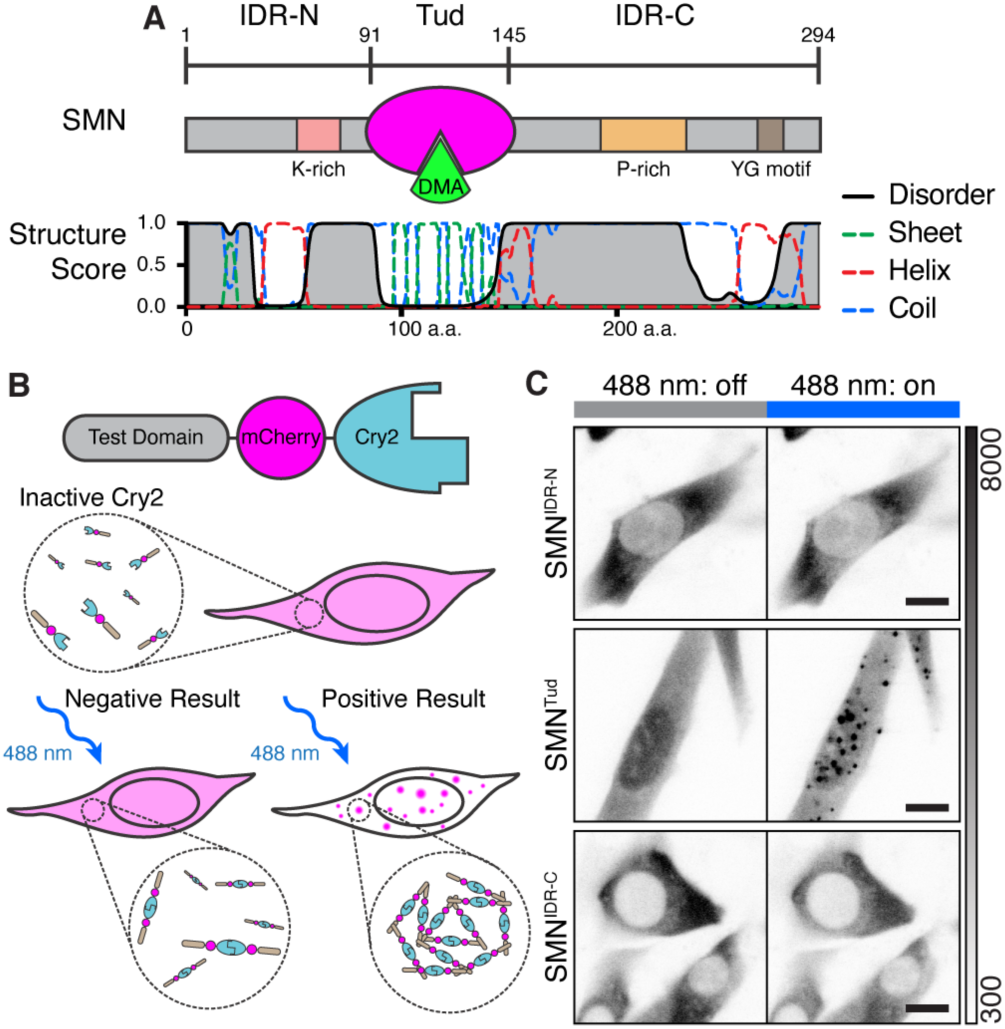
The multimerized SMN tudor domain forms condensates *in vivo*. A) Schematic representation of SMN domain architecture and accompanying secondary structure prediction. The tudor (Tud, magenta) domain binds DMA (green). Structure score is a unitless value for secondary structural properties predicted by the RaptorX algorithm. B) Diagram of the Cry2 condensation assay. Cry2 dimerizes upon blue light activation (488 nm); without added molecular interaction contributed by the test domain, condensation will not occur (Shin et al., 2017). If the test domain provides interactions to increase valency, condensation is observed as mCherry fluorescent foci. C) Micrographs of live cells undergoing blue light activation of Cry2 (180 s, blue bar). Grayscale bar given in analog-digital units. Scale bar = 10 μm.

Unexpectedly, SMN’s IDRs did not cluster, whereas the tudor domain (SMN^Tud^) formed prominent clusters throughout the cell upon light activation (Figure 1C; Figure S1B). These clusters were heterogenous and dynamic; when photobleached, they showed 42 - 100% recovery with time constants of 30 ± 22 s (mean ± SD, n=10) (Figure 2A). When Cry2 was allowed to deactivate after cluster formation, SMN^Tud^ clusters dissipated within a few minutes (Figure 2B). Cluster fusion was infrequent but observable (Figure 2C). Together these results show that SMN^Tud^ clusters rapidly reorganize their internal molecular composition, consistent with liquid-like condensation rather than stable aggregates. We conclude that SMN^Tud^ is sufficient for the formation of dynamic condensates upon light-induced multimerization.

**Figure 2.**
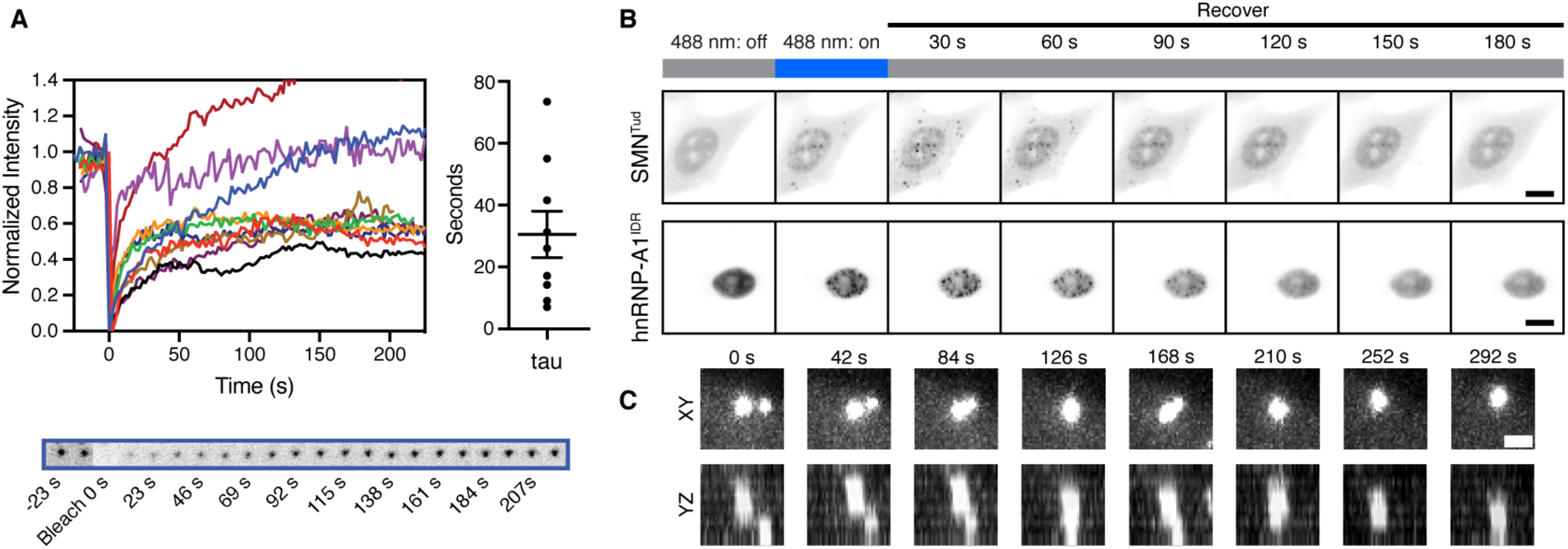
SMN^Tud^ dynamic clusters behave like liquid condensates. A) Fluorescence recovery after photobleaching for ten SMN^Tud^ condensed droplets with corresponding tau values from single exponential fits. Example images from blue trace provided below. B) Examples of condensate dissipation after Cry2 inactivation in live cells. Blue light pulse is 30 seconds. Scale bars = 10 μm. C) Example of a fusion event of two condensates. Scale bar = 2 μm.

This finding was striking because tudor domains are small (60 amino acids) and structured, rather than disordered and repetitive in sequence like FUS^IDR^ and hnRNP-A1^IDR^. We therefore asked whether condensate formation by SMN^Tud^ depends on the presence of its ligands, which are DMA modified proteins. To do so, we developed an image analysis method to quantify condensate formation compared to protein expression level. Our clustering metric is based on variance of intensity for each pixel throughout the movie recorded during Cry2 activation (Figure S2A). Correcting for the noise model of the camera and artefactual effects on variance, we obtained a clustering metric for each cell by averaging all pixels within segmented cell-by-cell masks and plotted it against the cell’s mean mCherry fluorescence (STAR Methods, Figure S2B&C). We determined a statistical threshold (Mann-Whitney U test) for the clustering metric of each cell relative to the mCherry-Cry2 negative control and found it to be at a clustering metric of approximately 49.44 ADU (Figure S2C).

DMA modifications can either be asymmetric or symmetric (aDMA or sDMA), and each is produced by separate sets of methyltransferase enzymes (Figure S2D) (Branscombe et al., 2001; Tang et al., 2000). The SMN tudor domain has a higher affinity for sDMA but recognizes both sDMA and aDMA (K_d_ = 0.476 mM and Kd = 1.025 mM respectively) (Tripsianes et al., 2011). Thus we chose two small-molecule inhibitors, MS-023 and EPZ015666 to deplete DMA modifications in NIH-3T3 cells (Figure S2E). MS-023 is a specific inhibitor of type I methyltransferases that synthesize aDMA (Eram et al., 2016). EPZ015666 is a specific inhibitor of protein methyltransferase 5, which synthesizes most sDMA (Chan-Penebre et al., 2015). We found that prolonged DMA inhibition substantially reduced the number of cells that form condensates above our significance threshold. (Figure 3A-C; Figure S2F). From this we conclude that condensation of SMN^Tud^ depends on the availability of endogenous DMA ligands.

**Figure 3.**
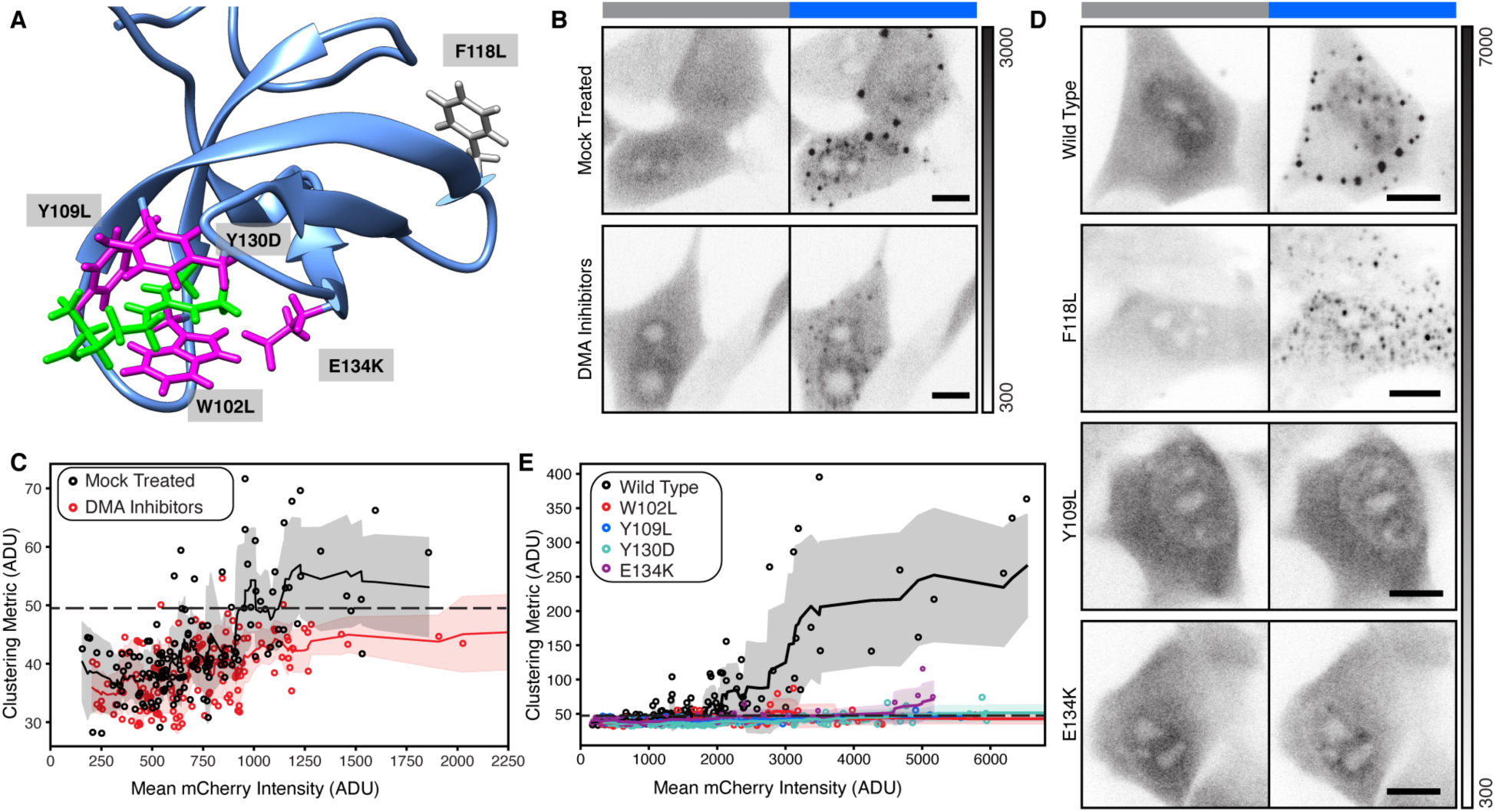
Formation of condensates by SMN^Tud^ depends on binding to DMA. A) Published solution structure of the SMN tudor domain (blue) bound to sDMA (magenta). Three aromatic amino acids that make up the binding pocket and one associated with SMA are shown in green. An aromatic residue (F118L) not involved in DMA binding is shown in gray. PDB: 4A4E*(Tripsianes et al*., *2011)*. B & D) Micrographs of live cells with SMN^Tud^ either untreated or treated with DMA inhibitors (B), SMN^Tud^ wild type, a null mutation F118L, a mutation to the aromatic DMA binding cage Y109L, and an SMA-associated mutation E134K (D). Grayscale bar given in analog-digital units. Scale bars = 10 μm. C & E) Quantification of live cells untreated or untreated with DMA inhibitors (C), cells expressing wild type or mutant SMN^Tud^ (E). Mean mCherry intensity and the cluster metric are given in analog-digital units (ADU). Solid line with shading is a rolling mean and standard deviation of 10 points. Each point is one cell. Dashed line represents a significance threshold relative to mCherry-Cry2 where α = 0.2 for the Mann-Whitney U test.

To verify that the loss of condensation is due to specific recognition of DMA by SMN^Tud^, we tested a series of mutants in the optodroplet assay. We chose three individually mutated amino acids in the aromatic cage that accepts DMA (W102L, Y109L, & Y130D), a disease causing mutation associated with SMA (E134K) that also disrupts DMA binding, and an uninvolved phenylalanine as a control (F118L) (Figure 3A) (Buhler et al., 1999; Tripsianes et al., 2011). All four mutations that disrupt DMA-binding virtually eliminated condensation, but F118L did not (Figure 3D&E; Figure S3). From these combined data, we conclude that condensation of SMN^Tud^ depends on binding to DMA ligands.

Tudor domains, including the SMN tudor, are known to bind DMA on multiple targets (Liu et al., 2012; Tripsianes et al., 2011). Therefore, we began testing known interactors of SMN for co-condensation with SMN^Tud^. DMA modified Sm proteins in snRNPs are canonical interactors of SMN, making them an obvious first candidate (Friesen et al., 2001). No snRNP marker colocalized with SMN^Tud^ condensates, indicating that Sm proteins do not participate in optodroplet formation by SMN^Tud^ (Figure S4). In contrast, endogenous coilin – a scaffolding protein of Cajal bodies that is modified by DMA – was detected in SMN^Tud^ nuclear condensates (Figure 4A). These nuclear condensates recruit coilin from a soluble pool, because the NIH-3T3 cells used do not have canonical (0.5 – 1.0 µm) Cajal bodies, though they do express coilin. To determine if coilin association is essential for condensate formation by SMN^Tud^, *Coil* knockout mouse embryonic fibroblasts were transduced with the Cry2 construct, and SMN^Tud^ condensates still formed in response to blue light (Figure 4B). These observations confirm that SMN^Tud^ can recruit coilin, a DMA ligand, to condensates. We conclude that coilin is a client of SMN^Tud^ in the context of our assay: it partitions into SMN^Tud^ condensates but is not necessary for their formation.

**Figure 4.**
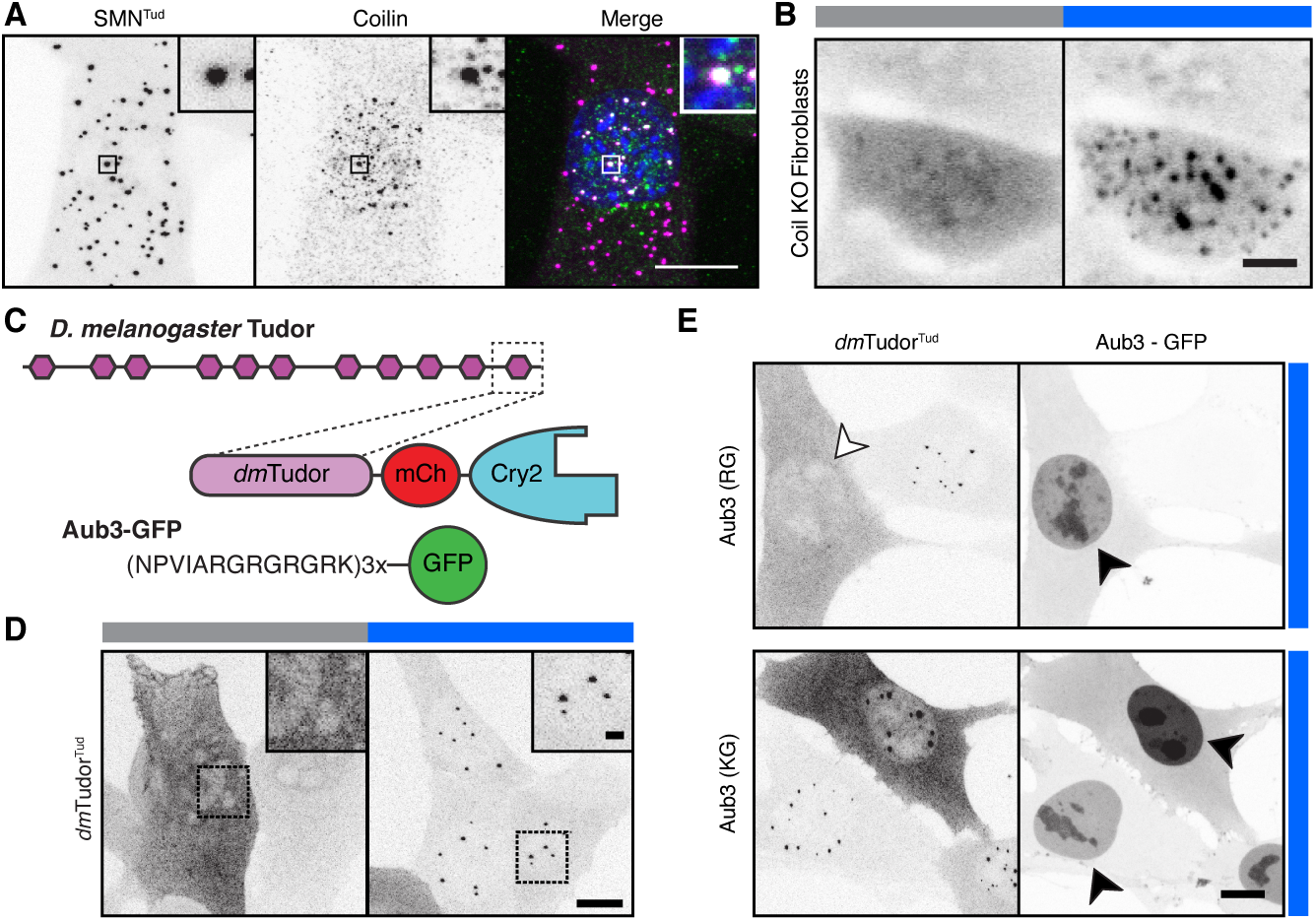
Methylated ligands partition into and compete for tudor domain binding sites. A) Light-activated and formaldehyde fixed cells expressing SMN^Tud^ (magenta) and stained for endogenous coilin (green), showing coilin is a client of SMN^Tud^. Hoechst staining shown in blue.B) Micrographs of Cry2-inactive (488 nm: off, grey) and Cry2-active (488 nm: on, blue) SMN^Tud^ in coilin knock-out mouse fibroblasts, showing coilin is not required for SMN^Tud^ condensation. C) Schematic of *D. melanogaster* Tudor, *dm*Tudor^Tud^ and its ligand Aub3-GFP (Liu et al., 2010). D) Fixed cells expressing only *dm*Tudor^Tud^ in Cry2-inactive and Cry2-active states, revealing condensation property of *dm*Tudor^Tud^. E) Fixed cells expressing *dm*Tudor^Tud^ and either Aub3-GFP or the non-binding, control peptide with R to K mutations (KG). Cry2 is active in all frames (blue). Condensates are inhibited by expression of intact Aub3-GFP (open arrowhead); both Aub3-GFP proteins concentrate in the nuclei (filled arrowheads) of transfected cells that lack condensates. Scale bars = 10 μm, inset scale bars = 2 μm.

To better understand the role of DMA ligands and to see if condensation of tudor domains is common, we turned to the eponymous protein of this domain family: *D. melanogaster* Tudor. The proteins Tudor, Aubergine, Vasa, and the oskar RNA make up the germ plasm condensate in flies (Trcek and Lehmann, 2019). DMA in the Aubergine N-terminus is required for germ plasm localization, and the binding between this region and the eleventh tudor domain of Tudor has been studied by co-crystallization (Liu et al., 2010). We generated a Cry2 construct with this tudor domain plus flanking sequence known to increase specificity (*dm*Tudor^Tud^) and a construct with three repeats of the Aubergine N-terminus fused to GFP (Aub_3_-GFP) (Figure 4C, Figure S5A). These constructs were introduced into NIH-3T3 cells. *Dm*Tudor^Tud^ was able to form condensates in the nucleus of NIH-3T3 cells without the expression of Aubergine showing that endogenous mouse ligands suffice (Figure 4D). Co-expression with Aub_3_-GFP, predominantly modified by aDMA, resulted in competition with the endogenous protein and blocked condensate formation (Figure 4E, Figure S5C). When lysine was substituted for arginine in Aub_3_-GFP, condensation was restored, confirming the necessity of the DMA-tudor interaction. The endogenous proteins that allowed *dm*Tudor^Tud^ to condense must be nuclear, because no condensation effect was seen in the cytoplasm. The recognition of a mouse protein by *dm*Tudor^Tud^ likely reflects the documented promiscuity of tudor domain recognition of DMA targets (Liu et al., 2012).

Given two examples of tudor domains that form condensates from two species, we asked if this property could be more broadly generalized. The human proteome has fifty-five annotated tudor domains from twenty-eight different proteins. Because each tudor domain binds a discrete set of DMA ligands due to amino acids surrounding the modified arginine, the condensation property observed for SMN^Tud^ could be unique (Liu et al., 2012; Tripsianes et al., 2011). Alternatively, the tendency to condense could be shared with other tudor domains. To test for generality, we selected a panel of twelve domains from nine different human proteins from diverse biochemical processes (Figure 5A). The twelve selected have the four aromatic amino acids required to form the binding pocket for DMA (Figure S5D). Of these domains, we found evidence for six that form condensates (Figure 5B, Figure S5E). As constructed, the assay is best suited to rule-in rather than rule-out domains, so we employed the clustering metric in live cells to assess whether expression level alone dictates whether a tudor domain makes condensates (Figure S5A). We compared SMN^Tud^ and Spf30^Tud^ and found that Spf30^Tud^ did not condense across a range of concentrations in spite of sequence, structural and interactome similarity to SMN^Tud^ (Figure S5E) (Tripsianes et al., 2011). This difference suggests that the ability of any tudor domain to mediate condensation depends on the availability, expression level, and methylation status of suitable partner molecules.

**Figure 5.**
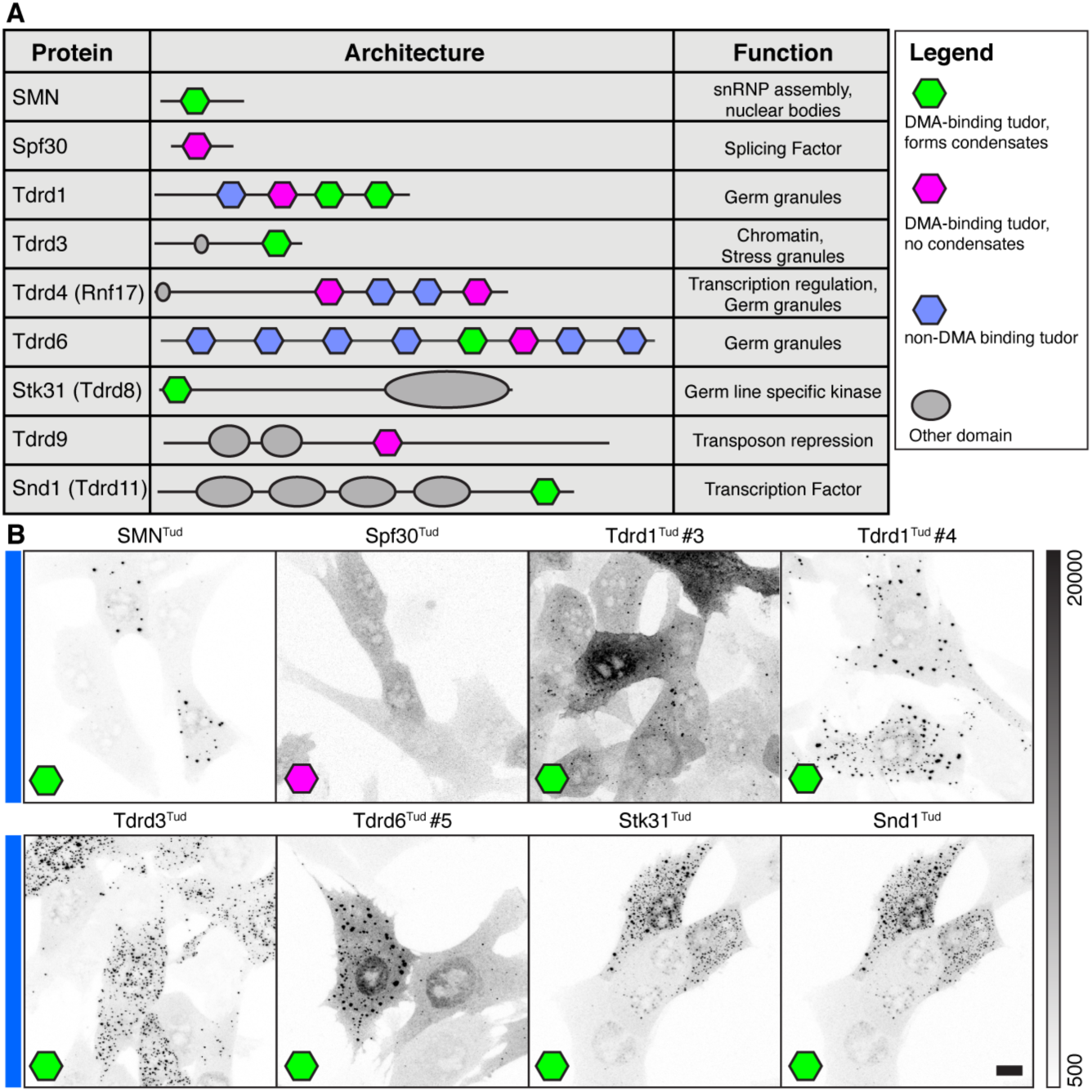
Condensation is a shared property of multiple human tudor domains. A) Table of proteins and schematics of human tudor domain proteins tested for condensation, with tudor domains containing either an intact binding site for DMA that form condensates (green), DMA binding tudor domains that do not form condensates (pink), or tudor domains lacking the DMA binding pocket (blue). Domain architecture and function correspond to Uniprot annotations.B) Fixed NIH-3T3 cells expressing Tudor-Cry2 constructs under Cry2-active conditions, corresponding to eight of the tudor domains above (see Figure S5 for more details). Grayscale bar given in analog-digital units. Scale bar = 10 μm.

To better understand how DMA-tudor interactions affect endogenous MLOs, we asked what specific effects arginine dimethylation has on the Cajal body. We treated HeLa cells with the specific inhibitors of DMA synthesis, MS-023 and EPZ015666, on HeLa cells and monitored their effects on Cajal bodies. Typically, HeLa cell Cajal bodies contain partially overlapping coilin and SMN domains (Figure 6A). The direction of these shifts is random but consistently observed ruling out chromatic aberration as their source (Figure S6A). Inhibition by both inhibitors results in the disassembly of Cajal bodies where residual coilin puncta lack the trimethylguanosine snRNP marker, in agreement with previous findings using a non-specific methylation inhibitor (Figure 6B; Figure S6B,C) (Hebert et al., 2002). We infer that the SMN bodies retained during DMA inhibition are gems, which are not dependent on DMA (Young et al., 2001). Thus, the integrity of the Cajal body nuclear organelle depends on modification of arginine, presumably on coilin.

**Figure 6.**
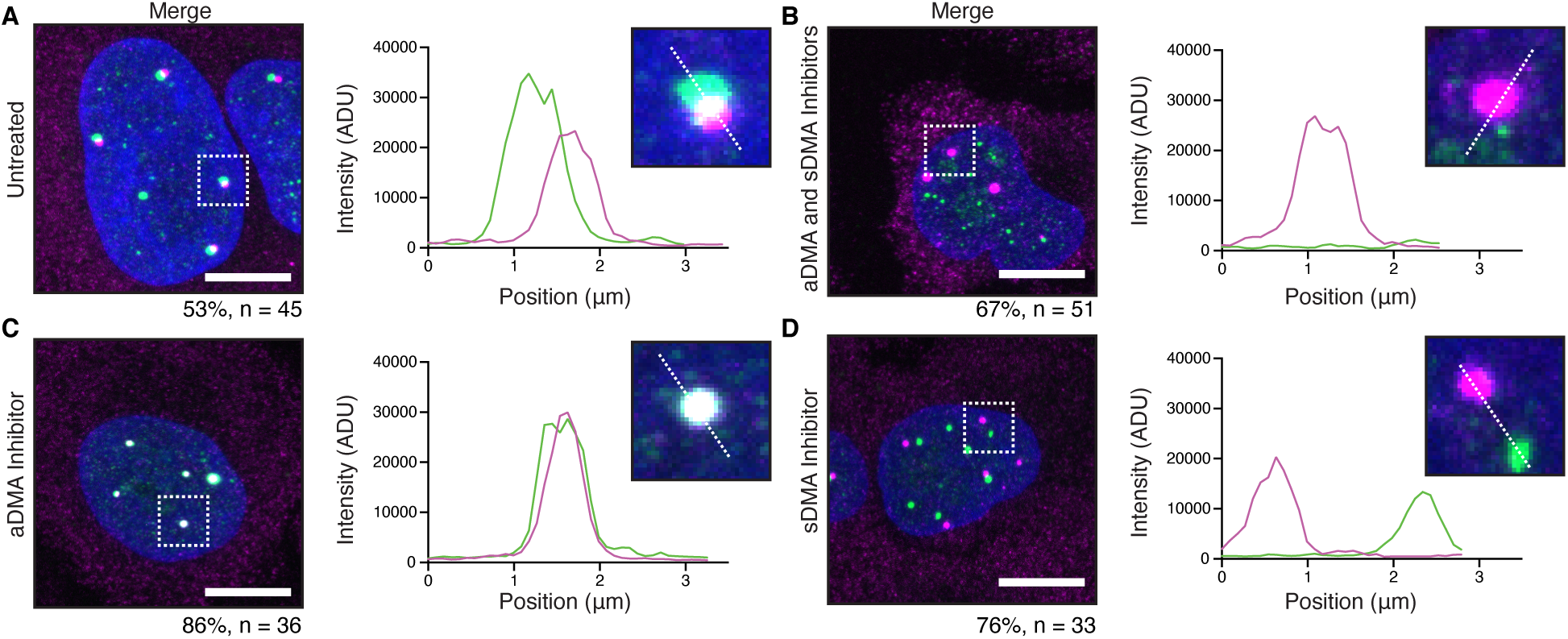
Specificity of nuclear body composition depends on DMA modification. Wild-type HeLa cells that are A) untreated, B) treated with MS023 and EPZ015666, C) treated with only MS023, or D) treated with only EPZ015666, and stained for SMN (magenta) and coilin (green) with accompanying inset and line profile plots. Cajal bodies and gems completely overlap when only sDMA is present; they are completely separate when only aDMA is present. Percentage of cells with the displayed phenotype out of total given below image. Scale bars = 10 μm.

Treatment of cells with individual inhibitors revealed an important requirement for the organization of Cajal bodies, gems, and their specificity. Inhibition of aDMA synthesis merges coilin and SMN into one fully overlapping body (Figure 6C). In contrast, inhibition of sDMA synthesis induces the complete separation of the coilin and SMN domains into two distinct bodies (Figure 6D). Importantly, trimethylguanosine continues to colocalize with coilin in both conditions, indicating snRNPs are still present with coilin in Cajal bodies with both individual inhibitors (Figure S6E,F). Thus, the Cajal body is dependent on arginine methylation overall, and symmetrical dimethylarginine – the highest affinity SMN tudor domain ligand (Tripsianes et al., 2011)– is required for the merging of Cajal bodies and gems into a single nuclear body. We suggest that normal levels of sDMA and aDMA appear to promote “docking” of the Cajal body to gems. These two organelles can either share components or retain unique identities based on the specific form of arginine methylation. Taken together, these observations demonstrate that the DMA-tudor module controls the compositional specificity of these nuclear MLOs.

## Discussion

Two key concepts emerge from this study. First, globular tudor domains promote condensation *in vivo*, by binding specifically to ligands bearing the modified amino acid dimethylarginine or DMA. We name this active unit the “DMA-tudor module” (Figure 7). Second, the particular constellation of DMA-tudor modules comprising aDMA and sDMA ligands determines the integrity of endogenous membraneless organelles (MLOs) exemplified by Cajal bodies and gems. We show that Cajal bodies require aDMA and sDMA and that DMA determines whether Cajal bodies and gems associate with one another. It was previously assumed that these MLOs mix their contents (Hebert and Matera, 2000). Surprisingly, our observations indicate these MLOs remain distinct and instead “dock” with one another when aDMA and sDMA are both present (Figure 7). Though Cajal bodies are often assumed to form by biomolecular condensation (Corbet and Parker, 2020; Hyman et al., 2014; Shin and Brangwynne, 2017). To our knowledge, this study provides the first mechanistic insights into how condensation contributes to the assembly and composition of this MLO. Moreover, the docking principle reveals rules governing the specificity of MLO composition. We provide evidence that additional DMA-tudor modules involving tudor domain-containing proteins with vastly different functions have the capacity to mediate condensation in other contexts, such as chromatin and germ plasm.

**Figure 7.**
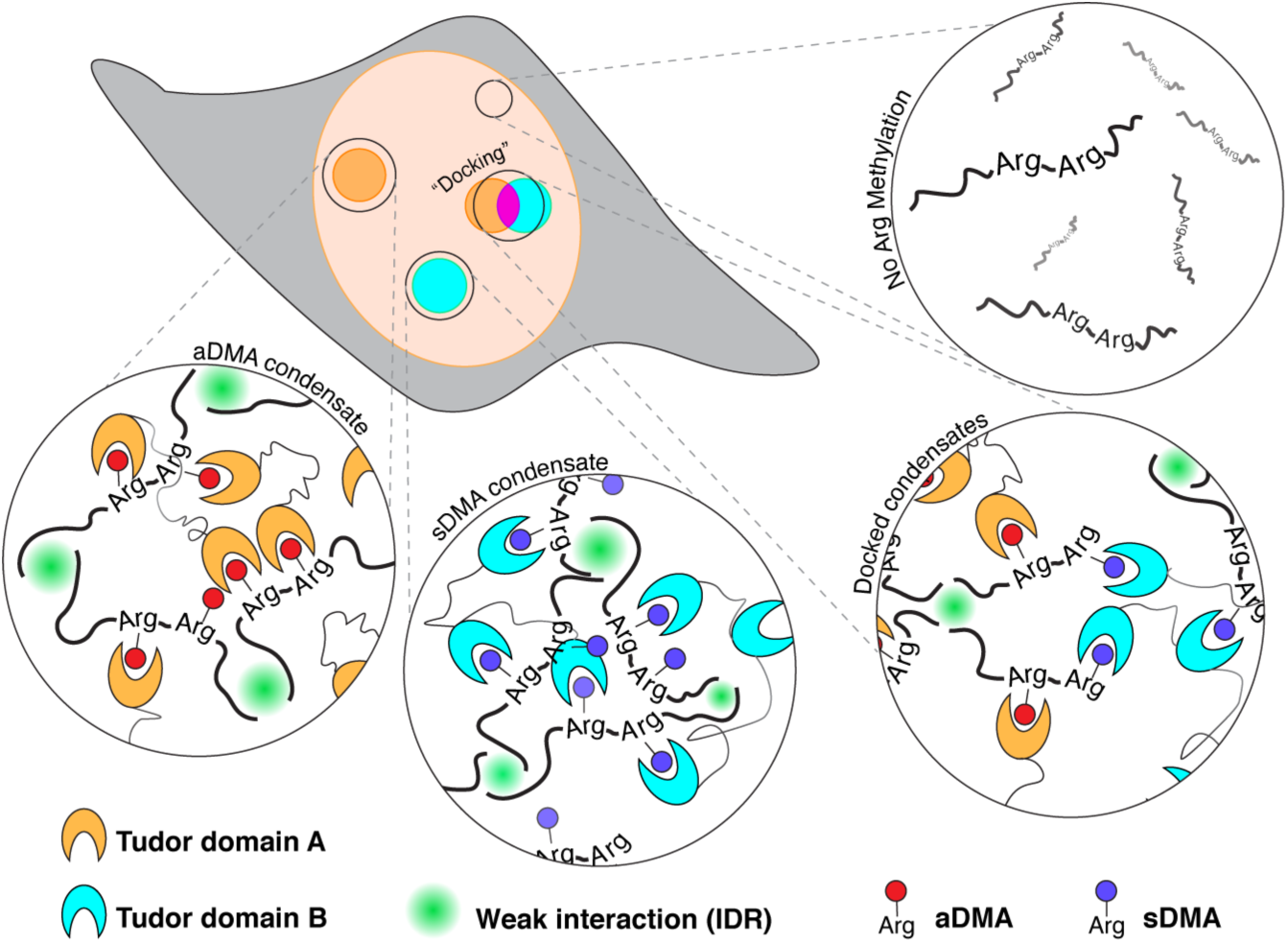
Proposed model for DMA-tudor module driven MLO formation. Working model of DMA modifications bound by tudor domains and how these DMA-tudor modules control the composition of MLOs. Unmodified arginine does not support assembly, while aDMA (red), and sDMA (blue) allow for assembly with tudor domain proteins (cyan and orange) in *trans*. If a protein bears both modifications, the two distinct bodies may “dock” with one another.

The DMA-tudor module is set apart from other specific interactions in condensates because it is based on a post-translational modification. In this way, it attains switch-like properties akin to the module formed by SH2 domains and phospho-tyrosine in Nck actin signaling condensates. However, DMA-tudor condensates are not nucleated on a two-dimensional lipid membrane and do not change the charge of the modified amino acid (Banjade et al., 2015). Nuclear MLOs like nucleoli and Cajal bodies are dynamic, disassembling during mitosis or cellular stress and reassembling quickly thereafter (Boulon et al., 2010; Strzelecka et al., 2010a). More broadly, remodeling of condensates in response to cell signaling has been proposed as a key mechanism of transcriptional via the phosphorylation of the RNA polymerase II C-terminal domain (Guo et al., 2019; Kwon et al., 2013). The modification of arginine has already been proposed as a way by which cells may modulate condensation by IDRs in *cis* (Nott et al., 2015; Ryan et al., 2018). We suggest that condensation regulation by DMA is more complicated than simply changing the chemical properties of the modified residues. Rather, DMA recruits the interaction of tudor domain proteins in *trans*. One demethylase for aDMA has been identified, making the modification reversible and potentially dynamic (Chang et al., 2007). Much is still unknown about how methyltransferases either differentiate between - or compete for – substrates for aDMA or sDMA modification.

Though we focused on tudor domains most likely to bind DMA, the tudor superfamily contains a spectrum of domains that bind other ligands or could bind DMA in a non-canonical manner (Chen et al., 2011). How DMA acts to regulate many condensates *in vivo* is unexplored. DMA modifications have been found on a vast and growing list of proteins including histones, RNA polymerase II, and G3BP1 (Chitiprolu et al., 2018; Tsai et al., 2016; Zhao et al., 2016). These proteins are associated with chromatin, transcription, and stress granules, respectively. Notably, all are systems reported to act as condensates *in vivo*, raising the possibility that tudor domain proteins and DMA play regulatory roles in their assembly and/or disassembly (Gibson et al., 2019; Guo et al., 2019; Molliex et al., 2015). The coactivator Tdrd3 uses its tudor domain to read aDMA modifications on core histone tails (H3R17me2a and H4R3me2a) and the CTD of RNA polymerase II at transcription start sites (Yang et al., 2010; Yang et al., 2014). During cellular stress, Tdrd3’s tudor domain targets the protein to stress granules (Goulet et al., 2008). Moreover, other tudor-domain regulators recognize DMA on histone tails (Lu and Wang, 2013). Based on our findings, we speculate that these activities may involve condensation by these tudor domains, which could also behave in a switch-like manner depending on the methylation state of the histone tails.

The Cry2 assay, combined here with analysis that relates condensation to protein expression level, reveals that a modest increase in valency through light-activated dimerization of tudor domains is enough to induce condensation (Bugaj et al., 2013; Shin et al., 2017). It is therefore striking that tudor domain proteins can have eight or more tudor domains not counting other interaction modules (Gan et al., 2019). The inherent multivalency of proteins like Tdrd4 and Tdrd6 implies a mechanism for germ line condensate formation, supported by our study of the original “Tudor” protein from *D. melanogaster*. Modified proteins often have multiple modified residues, offering avidity as an explanation for how relatively low affinity individual interactions allow for assembly (Tripsianes et al., 2011). Moreover, we show nuclear localization-specific condensation by *dm*Tudor^Tud^ and a lack of DMA-containing snRNPs in SMN^Tud^ condensates. Although there are many gaps in our knowledge about tudor domains and their ligands, each tudor domain exerts selectivity over the ligands bound, and not every methylated ligand is in every condensate. Since condensation is dependent on the concentration of the active unit required to separate from the bulk phase, the concentration of tudor domains and their available ligands likely determine when and whether condensates form (Elbaum-Garfinkle et al., 2015; Li et al., 2012; Pak et al., 2016). Thus, our data should be viewed as ruling-in certain tudor domains for condensation at the concentrations expressed stably in NIH-3T3 cells. Others may form condensates at higher concentrations, or in the presence of other ligands. Taken together, the DMA-tudor module has the properties of a highly versatile mechanism for cellular condensation.

At a minimum, two developmentally important MLOs – the Cajal body and germ plasm – are controlled by the DMA modification (Hebert et al., 2002; Liu et al., 2010; Nott et al., 2015). Our findings explain how DMA modification is able to control the composition of the Cajal body and its relationship to gems. Prior studies had observed that coilin methylation is required for its interaction with SMN, and inhibition of all cellular methylation disrupts Cajal body assembly (Hebert et al., 2002). Our work connects the current understanding of condensation to the past observation that coilin must bear a methylated arginine-glycine motif to form a Cajal body (Hebert et al., 2001). Furthermore, we observe that aDMA marks still promote Cajal body formation but deplete SMN from the organelle entirely. We presume that an as-yet unidentified tudor domain protein may be involved. Conversely, sDMA promotes the fusion of SMN-rich gems with Cajal bodies, showing that the precise form of this post translational mark dictates the occupancy of the body. Unperturbed, HeLa cells appear to exist in a state where Cajal bodies and gems are “docked” to one another implying that an equilibrium between aDMA and sDMA exists in this cell line. Importantly, improper assembly of the Cajal body may occur as part of SMA pathology, as we observe that SMN E134K cannot form condensates. The nuclear role of SMN is still unclear, but our results point to its function as a platform for condensation of the Cajal body which is required for development in vertebrates (Raimer et al., 2017; Strzelecka et al., 2010b).

The rules governing specificity and composition of condensates have been intensely sought after. While IDRs contribute important interactions to the formation of condensates, it is becoming clear that specific interactions are also required to form endogenous organelles via phase separation (Wang et al., 2018). The base pairing and secondary structure of RNA has recently been shown to play an important role (Langdon et al., 2018; Van Treeck et al., 2018). *In vitro* studies have shown that molecular recognition between folded domains can promote occupancy in a condensate (Ditlev et al., 2018). Clearly, the prolific nature of both tudor domain-containing proteins and DMA-modified proteins leaves open many opportunities for this mechanism to play out throughout the cell. Our study reveals that DMA-tudor modules provide the requisite specificity for the formation of endogenous MLOs. We speculate that biomolecular condensation *in vivo* can be driven by specific interactions of each tudor domain with its DMA ligands. Furthermore, the switch-like binding changes induced by the symmetry of the installed methyl groups introduces versatility and regulation, which can in turn define the identity and specific composition of MLOs formed through the activity of DMA-tudor modules.

## Acknowledgments

We thank A. Mennone, C. Emanuel, D. Baddeley, D. Phizicky, and S. Prophet for technical assistance. We thank members of the Neugebauer Lab, J. Steitz, and J. Howard for discussion and feedback on this manuscript. NIH awards U01DA047734 (to J.B.), NINDS-F31NS105379 (to E.M.C), U01CA200147 TCPA-2017-Neugebauer (to K.M.N.). This work is solely the responsibility of the authors and does not necessarily represent the official views of the NIH.

## Author contributions

E.C. and K.M.N. designed the study. K.S. generated all constructs. E.C. prepared cell lines and performed experiments. E.C. and A.E.S.B. carried out image analysis. J.B. and K.M.N. supervised the study. All authors contributed to writing the manuscript.

## STAR Methods

### Resource Availability

#### Lead Contact

Further information and requests for resources and reagents should be directed to and will be fulfilled by the Lead Contact, Karla M. Neugebauer.

#### Materials Availability

Plasmids generated in this study are available upon request.

#### Data and Code Availability

Raw microscopy data made available at data.4dnucleome.org. Image analysis code available at github.com.

### Method Details

#### Plasmid Construction

The plasmid system for lentiviral generation included pHR_SFFV (generated by Wendell Lim, Addgene #79121), pMD2.G (generated by Didier Trono, Addgene # 12259), and pCMVR 8.74 (generated by Didier Trono, Addgene # 22036). pHR_SFFV containing the IDRs of FUS and hnRNP-A1 were gifts from Cliff Brangwynne (Princeton University). Constructs in Table S1 below were generated with the InFusion HD kit (Takara) by replacing the sequence upstream of mCherry-Cry2 in pHR_SFFV-hnRNPA1-mCherry-Cry2. Aubergine peptide constructs were generated by inserting sequence upstream of GFP in pLV-EGFP (generated by Pantelis Tsoulfas, Addgene #36083, modified by David Phizicky) and expressed from the eIF4a promoter. Point mutations to SMN^Tud^ were generated by site-directed mutagenesis.

#### Cell Culture and Cry2 line generation

HEK 293FT, NIH-3T3, HeLa, and *coil* ^*-/-*^*-*mouse embryonic fibroblasts (MEF) were cultured in DMEM supplemented with 10% FBS, penicillin, and streptomycin (Gibco). All cells were incubated at 37°C in a 5% CO_2_ atmosphere. Pools of cells stably expressing Cry2 constructs were generated by first transfecting either pHR_SFFV or pLV-EGFP with the desired insert, pMD2.G, and pCMV R8.74 into confluent HEK 293FT. Transfection was performed with the Fugene HD reagent (Promega). After 48-72 h, viral supernatant was harvested, filtered, and applied to NIH-3T3 or *coil*^*-/-*^*-*MEF(Tucker et al., 2001). For live cell imaging, NIH-3T3 cells were cultured in glass-bottomed 35 mm dishes (MatTek). Just before imaging, the media was replaced with Live Cell Imaging Solution (Invitrogen) supplemented with 20 mM glucose and 1 μg/mL Hoechst 33342 (Invitrogen).

#### Live Cell Imaging of Cry2 Condensates

All live cell imaging was carried out on a Bruker Opterra II Swept Field instrument. The system allows for simultaneous imaging of mCherry fluorescence with the 561 nm laser and field of view activation of Cry2 with the 488 nm laser. The instrument is equipped with a Photometrics Evolve 512 Delta EMCCD camera. A standard protocol was used to make all Cry2 measurements. A PlanApo 60X 1.2 NA water immersion objective (Nikon) and a slit aperture of 70 μm was used to image at four frames per second. Four-frame averaging resulted in an overall frame rate of one frame per second. The 561 nm laser was set to 80% of full power and the 488 nm was set to 25% of full power. Both were passed through a 20% neutral density filter. We measured power output at the objective to be 80-90 μW for 561 nm and 20-25 μW for 488 nm with a Thorlabs PM100D power meter and a S170C sensor.

For each field of view, 10 s of inactive Cry2 were recorded, followed by turning on the 488 nm laser to capture 180 s of active Cry2. At the end of the series, a reference image of nuclear Hoechst staining was collected using the 405 nm laser. A stage-top incubator was employed to maintain cells at 37°C throughout imaging.

#### Fluorescence Recovery after Photobleaching (FRAP)

FRAP measurements were taken using identical imaging conditions as above with the following modifications. The objective was changed to a PlanApo 100X 1.4 NA oil immersion objective (Nikon). Frame averaging was replaced with a maximum projection of 7-11 z-steps at each time point, 0.25 s exposure per frame. After 180 s of activation, a single condensate was photobleached using a pulse of 405 nm laser light. The background was subtracted from a bleached region of interest containing a condensate using an unclustered region of the same cell. The intensity was then normalized to the maximum and minimum intensity values. Each recovery curve was fit using a one-phase exponential to estimate the mobile fraction and characteristic recovery time (1/*e* recovery, tau).

#### DMA Inhibitor Treatment

DMA inhibition was achieved by either individual or simultaneous treatment with two drugs: MS-023 (Sigma) and EPZ015666 (Sigma) *(Chan-Penebre et al*., *2015; Eram et al*., *2016)*. Stock solutions were prepared in DMSO at 50 mM and all subsequent dilutions were done in cell media as described above. MS-023 was added to media at a final concentration of 1.0 μM, and EPZ015666 was added at a final concentration of 5.0 μM. After 48 h of drug treatment, imaging was carried out as above, with drug added to the same concentration in the imaging media.

#### Fixed cell imaging

Cells were grown on No. 1.5 coverslips (Zeiss) in six-well plates and fixed in 4% paraformaldehyde (Sigma), blocked in 3% Bovine Serum Albumin (Sigma) and permeabilized by 0.1% Triton X-100 (American Bioanalytical). If immunofluorescence was performed, staining was conducted in blocking buffer with the antibodies listed in Table S2. For Cry2 activation prior to fixation, we built a custom illumination array designed for six-well plates. Six blue LEDs (470 nm, 3.2 V, Digi-Key 150080BS75000) were arrayed to illuminate each well in a six-well plate for 5 min (Extended Data Fig. 10b). Illumination was maintained during the 10 min fixation step. Inactive Cry2 samples were maintained in the dark for 10 min and fixed in the same manner. Imaging was done on a Leica Sp8 Laser Scanning Confocal.

#### Western Blotting

Western blots were carried out by preparing whole cell lysate in 0.1% SDS, 0.5% sodium deoxycholate, 150 mM NaCl, 10 mM EDTA, and 10 mM Tris (pH 8.0). Lysates were denatured and reduced by adding NuPAGE LDS sample buffer (Thermo) and heating for 10 min at 85°C. Samples were run out on a 4-12% acrylamide Bis-Tris gel (Invitrogen) and transferred to 0.2 μm nitrocellulose membranes (BioRad). After blocking, membranes were stained with the antibodies listed in Table S2.

### Quantification and Statistical Analysis

#### Image Analysis: Segmentation

Quantification of condensate formation on a per-cell basis requires accurate segmentation of cells within a field of view. A cell profiler pipeline was used to process movies into cell masks. Binary nuclear masks were generated by segmenting the Hoechst image associated with each sequence. Binary cellular masks were generated by performing an Otsu threshold and watershed transform seeded by the nuclear masks on the mCherry signal from inactive Cry2 cells. Masks defined by automated segmentation were manually checked for accuracy; cells with substantial movement from the mask were excluded from analysis.

#### Image Analysis: Intensity correction

We use a temporal variance metric to quantify clustering without imposing a model on the time-dependence or shape, in an expression-level independent manner. Fluorophores moving into or out of a pixel will increase the temporal variance, substantially for pixels seeing large influxes of fluorophores. However, as photon-counting processes are subject to shot-noise, the temporal variance of a pixel will also increase with baseline intensity increases, which would confound comparison between cells of differing expression levels. This effect can be conveniently normalized using the temporal mean, since the variance and expectation value of a Poisson process are equal. Several effects which deviate our raw analog-digital counts from this model were therefore accounted for before calculating the clustering metric.

First, the per-pixel analog-digital offset of the camera was characterized and subtracted from all images. We often observe very bright spots appearing on single frames at random positions, which we interpret to be particles or cosmic rays impacting the EMCCD. Such large spikes in intensity can bias variance-based calculations, and because these spikes sometimes occur within cell masks, we filter them using a temporal sliding-window approach (Extended Data Fig.10a). We iterate through time and calculate the median-absolute deviation (MAD) at each *xy*- pixel within a 10-frame window, where the MAD of a sequence *k* is simply median(|*k* − median(*k*)|). Values within this window were identified as spikes if they were both larger than 15 times the MAD and were increases on the previous frame larger than 1500 ADU. Spikes were replaced with the median value of the window, then the window was advanced one frame and the process was repeated until the last frame was included in the window. MAD was employed to determine the variability within a window as it is a robust variability measure, and does not increase substantially due to the very spikes we are trying to filter, unlike the standard deviation or variance.

Finally, we performed a mean-normalization to remove effects of read-out laser intensity fluctuation, bleaching, etc., by multiplying frame *j* by a factor *β =* ⟨*I*(*t* = 0) ⟩_*x,y*_*/* ⟨*I*(*t*_*j*_)⟩_*x*_,_*y*_ where ⟨*k*⟩_*x*_ denotes an average of *k* taken over *x*.

#### Image Analysis: Clustering Metric

We calculated the temporal variance and average for each pixel during Cry2-activation in live cells. In order to arrive at a metric which is theoretically independent of baseline signal magnitude, and therefore expression levels, we normalized the temporal variance by the mean. This is a convenient normalization for data influenced by shot-noise because the variance of a Poisson process is equal to its mean. The clustering metric is given by

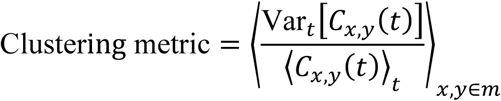

where *C*_*i,j*_*(t)* is the corrected intensity at pixel *x,y* as a function of time, and *m* denotes the set of *x,y* pixels within a mask. An intermediary variance-over-mean image can be generated where individual clusters are visible. Using fluorescence intensity as a proxy for concentration, the clustering metric can be plotted against baseline intensity so that comparisons between constructs may be made (Extended Data Fig. 2d,e).

The clustering metric for a homogeneous Poisson process should be unity, however our camera data is not only subject to shot noise, but also read noise and noise due to the stochastic nature of electron multiplication (Robbins and Hadwen, 2003). A more complete camera noise model may be necessary for comparison of results from different microscopes or camera settings.

#### Clustering classification

While the clustering metric allows for comparison between the degree of clustering for individual cells, the expression efficiency and/or sampling varies between constructs, complicating statistical analysis between them. To remove the influence of data which did not experience significant clustering, we performed a Mann-Whitney U test between the clustering metric distribution of the negative control (mCherry-Cry2) and each individual clustering metric from other distributions, classifying cells which reject the null hypothesis with *α* = 0.2 as cells which experience clustering. We then calculated the median clustering metric for a construct using only its cluster-positive cells and used that to compute differences in clustering magnitude.

#### Software

Images were prepared for figures and FRAP calculations were done using ImageJ (FIJI) version 2.0.0-rc-69/1.52p (Schindelin et al., 2012). Image segmentation was done using Cell Profiler version 3.1.9 (Carpenter et al., 2006). All other image analysis was done using PYME version 19.12.17 which is available at python-microscopy.org. The plugin used to compute clustering metrics can be found at github.com/bewersdorflab/quant-condensate. Structure prediction was performed on the RaptorX Property Prediction server using default parameters (Kallberg et al., 2012). Tudor domains were aligned by extracting domain annotations from Uniprot.org and aligning sequences with Clustal Omega using the default parameters (Sievers et al., 2011).

#### Image Display

All images within a figure are contrast matched, and an intensity grayscale or color bar is given for reference where applicable. All live cell images are micrographs from single z-slice movies except where otherwise noted. Images and quantification comparisons are only made between samples taken with identical microscope calibrations. Confocal images of fixed samples are maximum intensity projections of z-stacks.

**Figure S1. The Cry2 assay reveals the condensation properties of protein domains**.

a) Micrographs of live cells expressing mCherry-Cry2, hnRNP-A1^IDR^, and FUS^IDR^. mCherry does not cluster while the IDRs do. Results produced using the constructs and protocols from Shin, Y., et al.(Shin et al., 2017). b) Micrographs of live cells expressing fragments of SMN in fusion with mCherry-Cry2. Only the tudor domain forms clusters when Cry2 is active. Grayscale bars are in analog-digital units. Scale bar = 10 μm.

**Figure S2. Quantification of cluster formation reveals SMN**^**Tud**^ **dependence on DMA**

A) Live cells are recorded for 10 seconds without Cry2 activation and then 180 seconds with blue light which activates Cry2. The mCherry signal before activation allows for segmentation of individual cells and serves as a proxy for expression level. The per-pixel temporal variance and mean are computed for the Cry2-active period. These values are used to construct the per-pixel clustering metric (right). Per-cell values are calculated by averaging over each cell mask. Scale bar = 10 μm. Grayscale bar (left) indicates fluorescence intensity, color bar (right) indicates clustering metric, both given in analog-digital units (ADU). B) An example of a single frame spike in intensity filtered out using a MAD-based approach. Arrowhead indicates spike. Before filtering, a high intensity spot is seen inside the region of measurement for the clustering metric. After filtering, the spot is effectively removed. Scale bar = 10 μm. Each frame is a 1.0 s exposure. C) Clustering metrics for SMN^TUD^ and mCherry-Cry2 as a negative control. Solid lines and shading are a rolling mean and standard deviation of 10 points. Each point represents one cell. Dashed line represents a significance threshold relative to mCherry-Cry2 where α = 0.2 for the Mann-Whitney U test. D) Chemical structures of sDMA and aDMA. E) Whole cell lysates from NIH-3T3 cells from the same biological replicate treated for 48 h with MS-023, EPZ015666, or both, blotted for aDMA and sDMA. GAPDH blot is from separate lanes and shown next to both DMA blots. Numbered asterisks indicate 1) a band only reduced when treated with both drugs, 2) the major band reduced by MS-023, and 3) a band reduced by EPZ015666. F) Micrographs of live cells expressing SMN^Tud^ untreated and mock treated with DMA Inhibitors MS-023 and EPZ015666 for 48 h. Grayscale bar is in analog-digital units. Scale bar = 10 μm.

**Figure S3. Mutations that affect DMA binding eliminate SMN**^**Tud**^ **condensation**

A-E) Micrographs of live cells expressing A) SMN^Tud^ or SMN^Tud^ with point mutations B) W102L, C) Y109L, D) Y130D, and E) E134K that reduce binding to DMA *(Tripsianes et al*., *2011)*. F) Micrographs of live cells expressing F118L mutation to SMN^Tud^. Grayscale bars are in analog-digital units and applies to all images. Scale bars = 10 μm. G) Quantification of the condensation effect of SMN^Tud^ F118L compared to cells expressing mCherry-Cry2. Units are analog-digital units (ADU). Solid lines and shading are a rolling mean and standard deviation of 10 points. Each point represents one cell. Dashed line represents a significance threshold relative to mCherry-Cry2 where α = 0.2 for the Mann-Whitney U test.

**Figure S4. SMN**^**Tud**^ **condensates do not contain snRNPs**

Fixed cells with Cry2-active SMN^Tud^, which is always visualized by mCherry fluorescent signal. Condensates were stained for a) mCherry, b) sDMA using antibody SYM10, c) the Sm epitope with Y12, d) the trimethylguanosine 5′ cap, e) U1 snRNP-C, and f) U1-70k, both U1 snRNP specific proteins. In b) arrowheads indicate nuclear condensates that stain positively with SYM 10. Scale bar = 10 μm.

**Figure S5. Numerous human tudor domains form condensates**

A) Western blot of whole cell lysate from NIH-3T3 cells showing expression levels of Cry2 constructs as measured by mCherry staining. All Constructs include mCherry-Cry2. B) LED array used for activating Cry2 in cells plated in six-well plates with coverslips. C) Western blot of whole cell lysates of NIH-3T3 wild-type, expressing Aub3-GFP, or expressing Aub3-GFP with arginine mutated to lysine. Each panel shows the range of ∼25-40 kDa. D) Amino acid sequences of tudor domains aligned using Clustal Omega. Green highlights indicate aromatic residues in the DMA binding pocket as determined by structural studies. E) Fixed NIH-3T3 cells expressing Tudor- Cry2 constructs under Cry2-inactive and Cry2-active conditions. Color bar given in analog-digital units. Scale bar = 10 μm. F) Quantification of the condensation effect in live cells of SMN^Tud^ compared to Spf30^Tud^. Units are analog-digital units (ADU). Solid lines and shading are a rolling mean and standard deviation of 10 points. Each point represents one cell. Dashed line represents a significance threshold relative to mCherry-Cry2 where α = 0.2 for the Mann-Whitney U test.

**Figure S6. Specificity of Cajal body assembly is controlled by DMA**

a) Two biological replicates of untreated wild-type HeLa cells stained for SMN (magenta) and coilin (green). b) Untreated, c) MS023 and EPZ015666 treated, d) MS023 treated, and e) EPZ015666 treated HeLa cells stained for trimethylguanosine (magenta) and coilin (green). Arrowheads in insets indicate the coilin puncta shown in each line profile plot. Scale bar = 10 μm.

**Movie S1. mCherry-Cry2 does not cluster in the optodroplet assay**

NIH-3T3 cells expressing mCherry-Cry2. At time 0.0, blue light is turned on to activate Cry2. Time resolution, 1.0 frame per second. Scale bar = 10 µm.

**Movie S2. SMN**^**Tud**^ **forms clusters in the optodroplet assay**

NIH-3T3 cells expressing SMN^Tud^. At time 0.0, blue light is turned on to activate Cry2. Time resolution, 1.0 frame per second. Scale bar = 10 µm.

**Table S1. Details for constructs used in this study**

**Table S2. Details for antibodies used in this study**

