## Supplemental Figures & Info for "DMA-tudor interaction modules control the specificity of *in vivo* condensates"

**Figure S1. The Cry2 assay reveals the condensation properties of protein domains.**

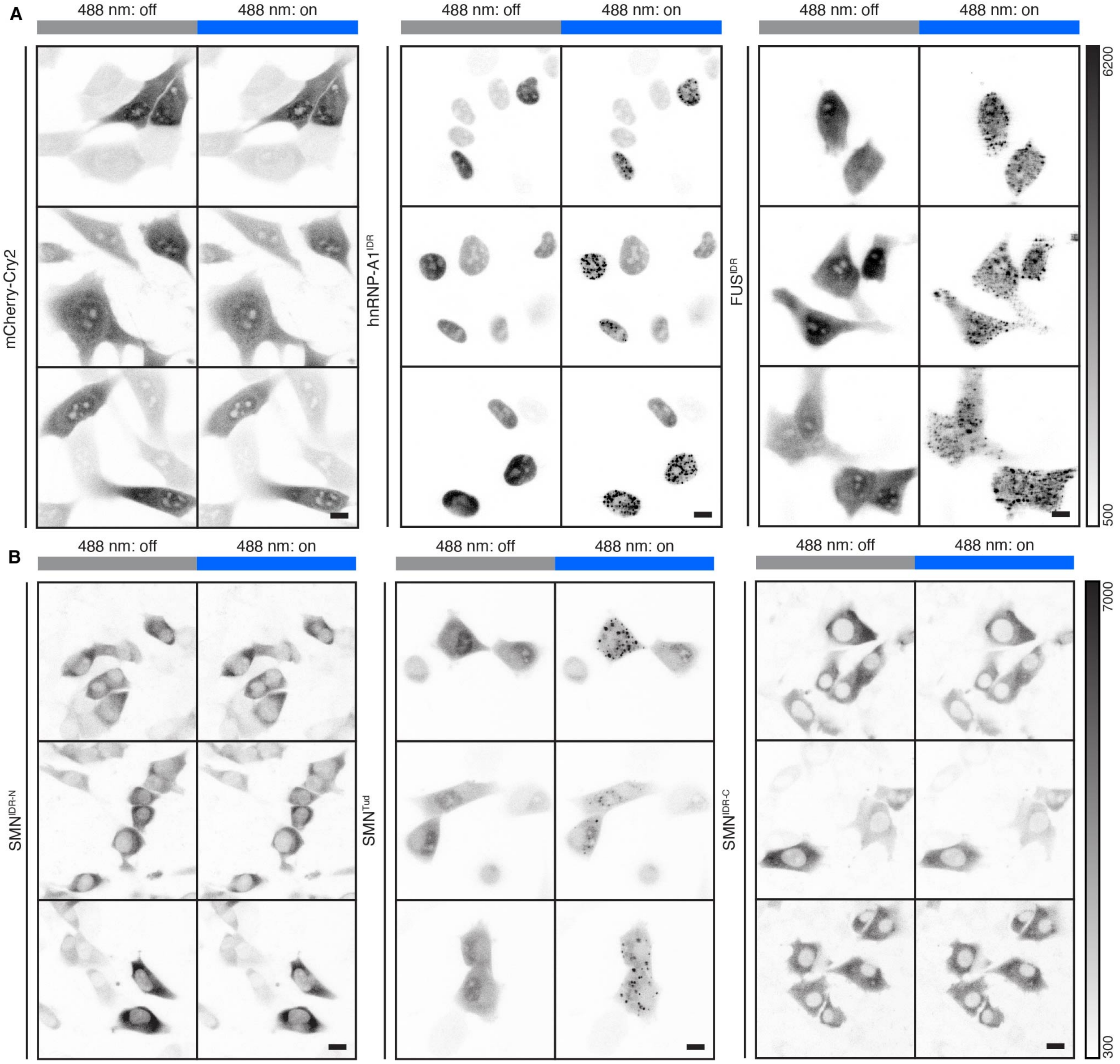


a) Micrographs of live cells expressing mCherry-Cry2, hnRNP-A1$^{IDR}$, and FUS$^{IDR}$. mCherry does not cluster while the IDRs do. Results produced using the constructs and protocols from Shin, Y., et al.(Shin et al., 2017). b) Micrographs of live cells expressing fragments of SMN in fusion with mCherry-Cry2. Only the tudor domain forms clusters when Cry2 is active. Grayscale bars are in analog-digital units. Scale bar = 10 μm.

**Figure S2. Quantification of cluster formation reveals $SMN^{Tud}$ dependence on DMA.**

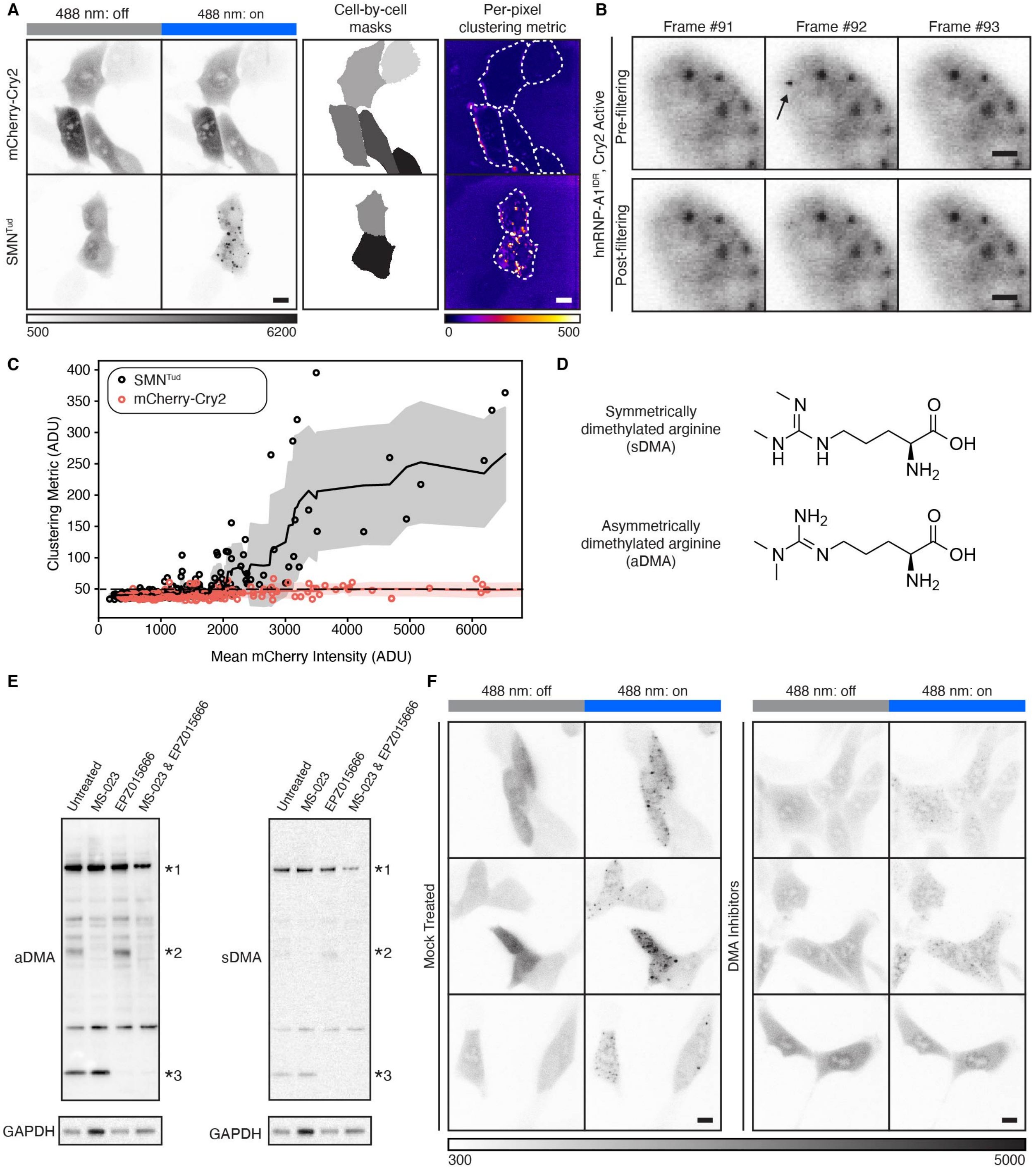


A) Live cells are recorded for 10 seconds without Cry2 activation and then 180 seconds with blue light which activates Cry2. The mCherry signal before activation allows for segmentation of individual cells and serves as a proxy for expression level. The per-pixel temporal variance and mean are computed for the Cry2-active period. These values are used to construct the per-pixel clustering metric (right). Per-cell values are calculated by averaging over each cell mask. Scale bar = 10 μm. Grayscale bar (left) indicates fluorescence intensity, color bar (right) indicates clustering metric, both given in analog-digital units (ADU). B) An example of a single frame spike in intensity filtered out using a MAD-based approach. Arrowhead indicates spike. Before filtering, a high intensity spot is seen inside the region of measurement for the clustering metric. After filtering, the spot is effectively removed. Scale bar = 10 μm. Each frame is a 1.0 s exposure. C) Clustering metrics for $SMN^{TUD}$ and mCherry-Cry2 as a negative control. Solid lines and shading are a rolling mean and standard deviation of 10 points. Each point represents one cell. Dashed line represents a significance threshold relative to mCherry-Cry2 where a = 0.2 for the Mann-Whitney U test. D) Chemical structures of sDMA and aDMA. E) Whole cell lysates from NIH-3T3 cells from the same biological replicate treated for 48 h with MS-023, EPZ015666, or both, blotted for aDMA and sDMA. GAPDH blot is from separate lanes and shown next to both DMA blots. Numbered asterisks indicate 1) a band only reduced when treated with both drugs, 2) the major band reduced by MS-023, and 3) a band reduced by EPZ015666. F) Micrographs of live cells expressing $SMN^{Tud}$ untreated and mock treated with DMA Inhibitors MS-023 and EPZ015666 for 48 h. Grayscale bar is in analog-digital units. Scale bar = 10 μm.

**Figure S3. Mutations that affect DMA binding eliminate SMN$^{Tud}$ condensation.**

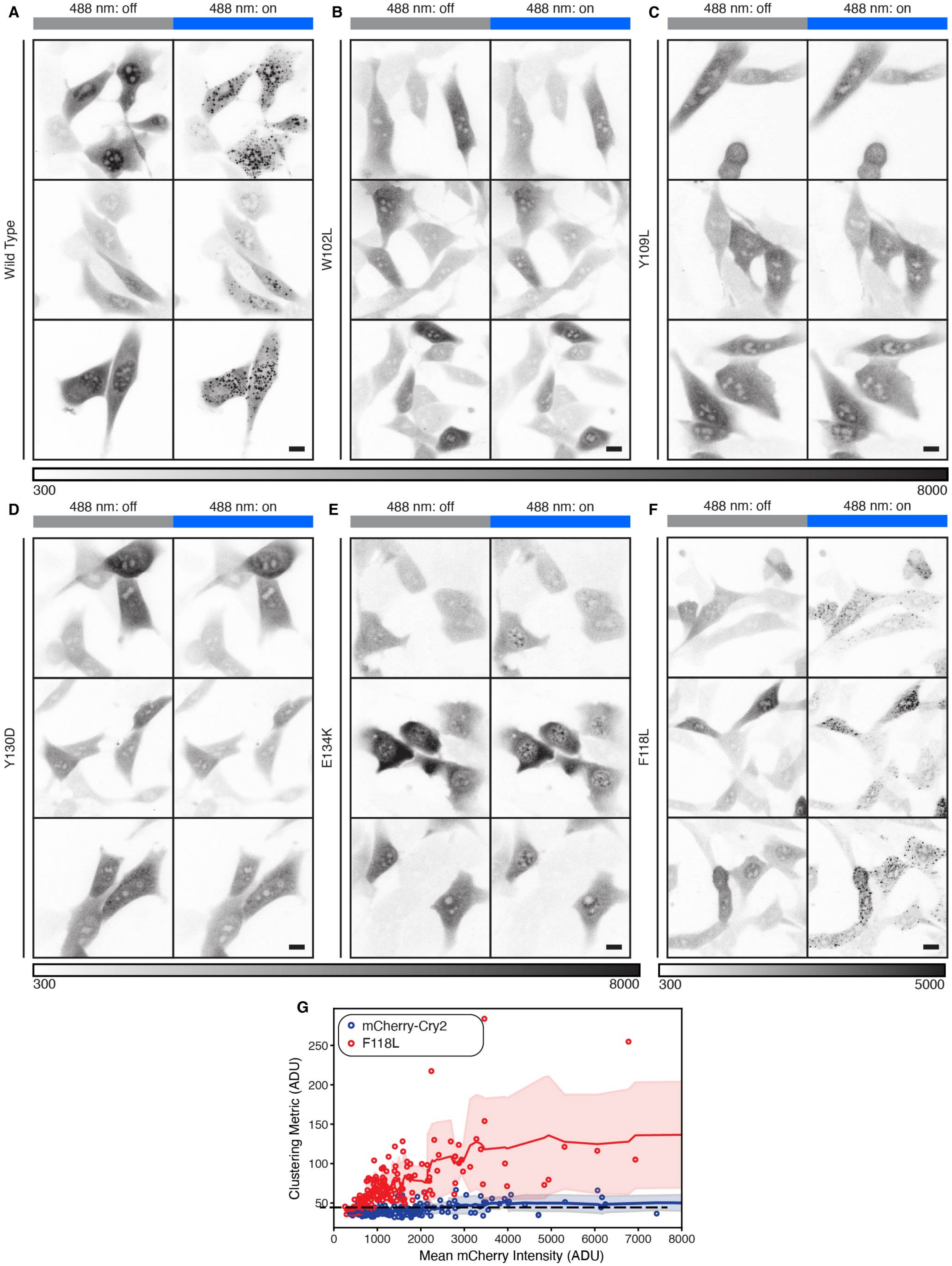

A-E) Micrographs of live cells expressing A) $SMN^{Tud}$ or $SMN^{Tud}$ with point mutations B) W102L, C) Y109L, D) Y130D, and E) E134K that reduce binding to DMA *(Tripsianes et al., 2011).* F) Micrographs of live cells expressing F118L mutation to $SMN^{Tud}$. Grayscale bars are in analog-digital units and applies to all images. Scale bars = 10 µm. G) Quantification of the condensation effect of $SMN^{Tud}$ F118L compared to cells expressing mCherry-Cry2. Units are analog-digital units (ADU). Solid lines and shading are a rolling mean and standard deviation of 10 points. Each point represents one cell. Dashed line represents a significance threshold relative to mCherry-Cry2 where a = 0.2 for the Mann-Whitney U test.

**Figure S4. SMN$^{Tud}$ condensates do not contain snRNPs.**

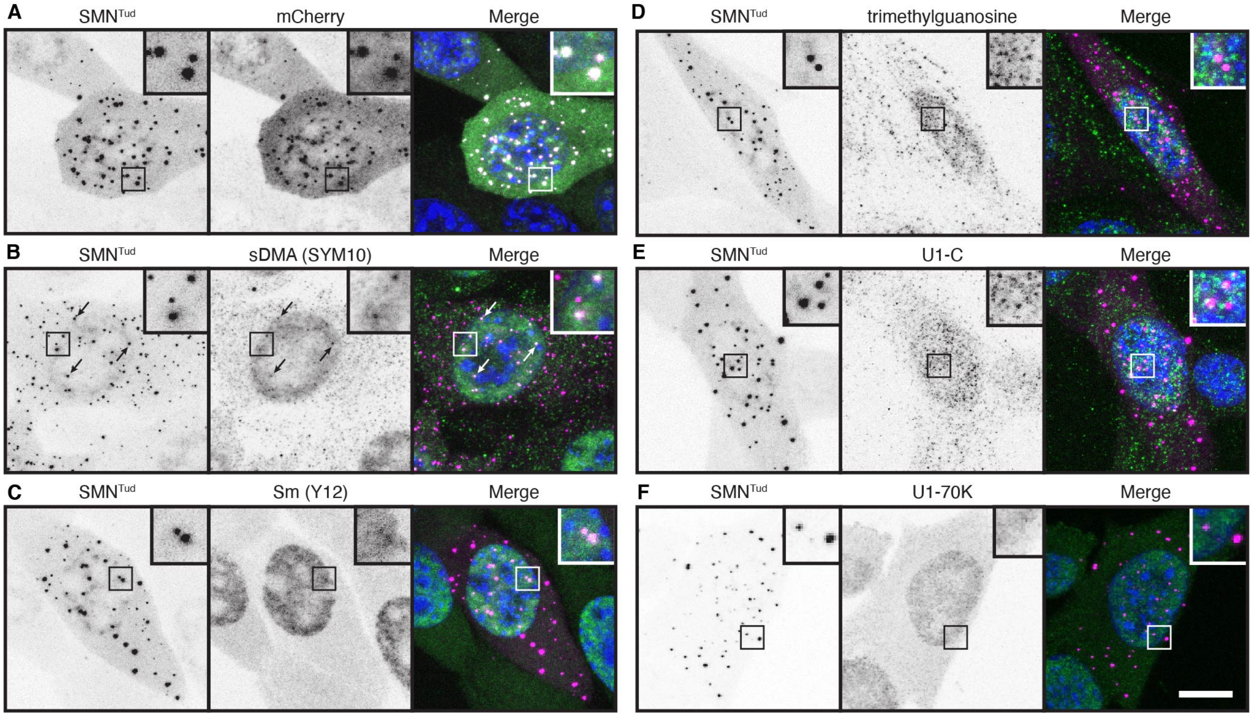


Fixed cells with Cry2-active SMN$^{Tud}$, which is always visualized by mCherry fluorescent signal. Condensates were stained for a) mCherry, b) sDMA using antibody SYM10, c) the Sm epitope with Y12, d) the trimethylguanosine 5′ cap, e) U1 snRNP-C, and f) U1-70k, both U1 snRNP specific proteins. In b) arrowheads indicate nuclear condensates that stain positively with SYM 10. Scale bar = 10 μm.

**Figure S5. Numerous human tudor domains form condensates.**

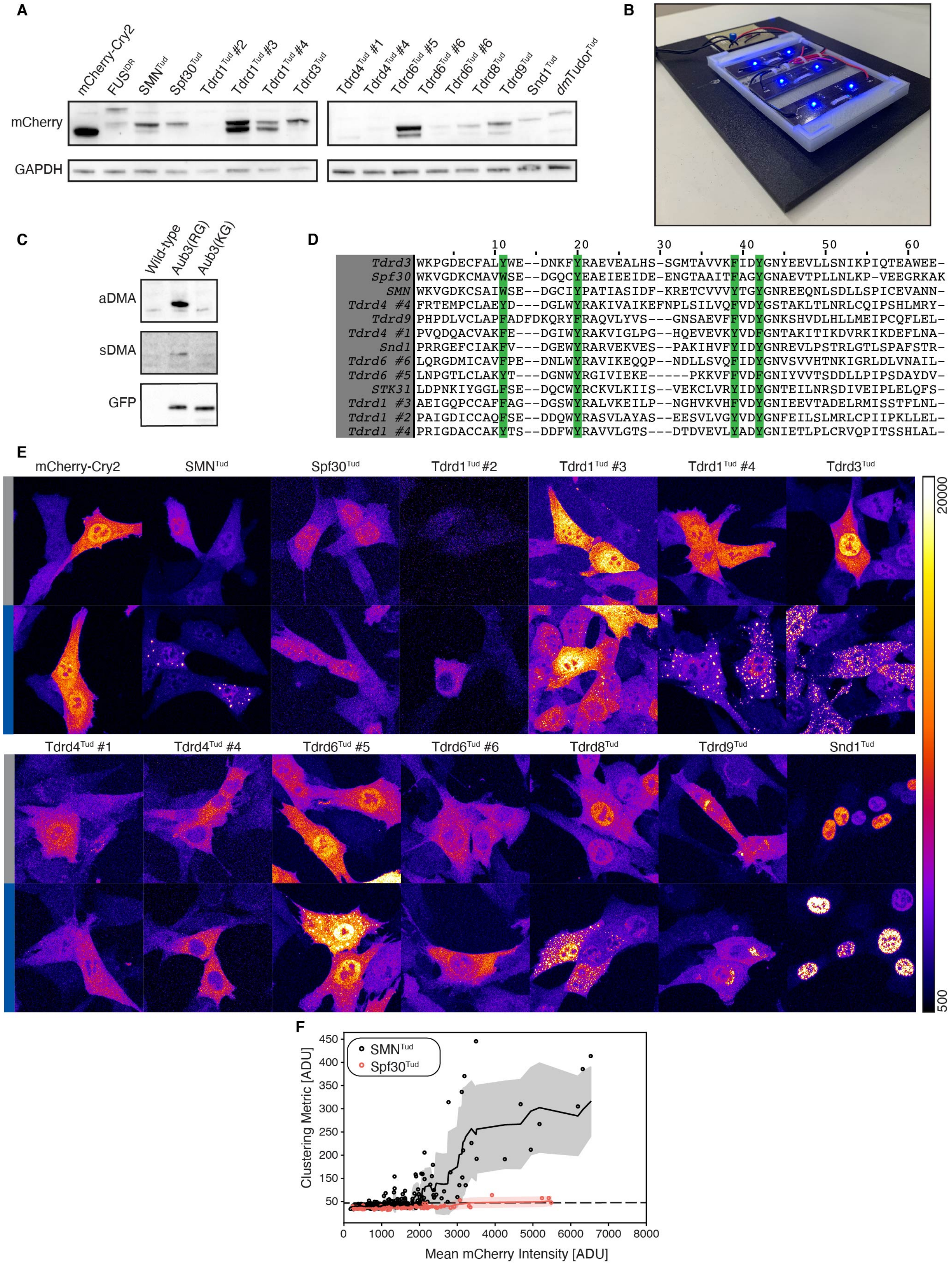

A) Western blot of whole cell lysate from NIH-3T3 cells showing expression levels of Cry2 constructs as measured by mCherry staining. All Constructs include mCherry-Cry2. B) LED array used for activating Cry2 in cells plated in six-well plates with coverslips. C) Western blot of whole cell lysates of NIH-3T3 wild-type, expressing Aub3-GFP, or expressing Aub3-GFP with arginine mutated to lysine. Each panel shows the range of ~25-40 kDa. D) Amino acid sequences of tudor domains aligned using Clustal Omega. Green highlights indicate aromatic residues in the DMA binding pocket as determined by structural studies. E) Fixed NIH-3T3 cells expressing Tudor-Cry2 constructs under Cry2-inactive and Cry2-active conditions. Color bar given in analog-digital units. Scale bar = 10 μm. F) Quantification of the condensation effect in live cells of $SMN^{Tud}$ compared to $Spf30^{Tud}$. Units are analog-digital units (ADU). Solid lines and shading are a rolling mean and standard deviation of 10 points. Each point represents one cell. Dashed line represents a significance threshold relative to mCherry-Cry2 where a = 0.2 for the Mann-Whitney U test.

**Figure S6. Specificity of Cajal body assembly is controlled by DMA.**

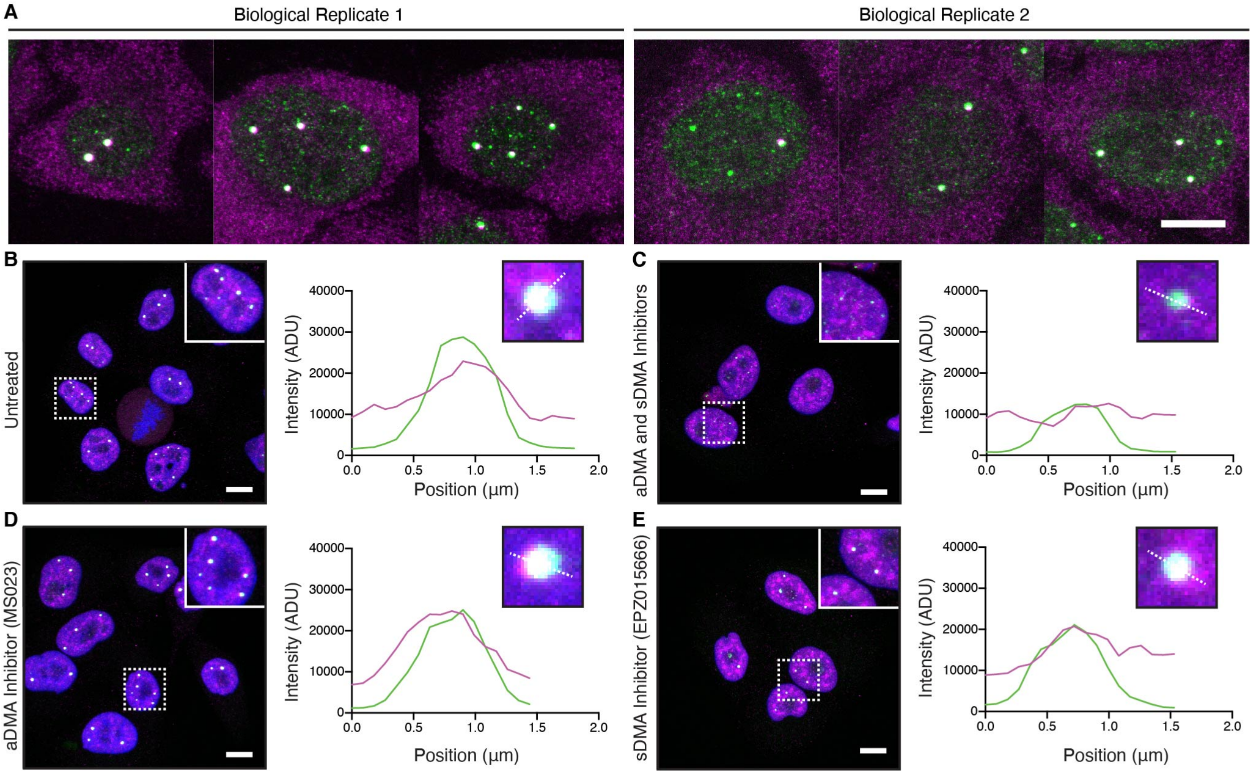


a) Two biological replicates of untreated wild-type HeLa cells stained for SMN (magenta) and coilin (green). b) Untreated, c) MS023 and EPZ015666 treated, d) MS023 treated, and e) EPZ015666 treated HeLa cells stained for trimethylguanosine (magenta) and coilin (green). Arrowheads in insets indicate the coilin puncta shown in each line profile plot. Scale bar = 10 µm.

**Movie S1. mCherry-Cry2 does not cluster in the optodroplet assay.**

NIH-3T3 cells expressing mCherry-Cry2. At time 0.0, blue light is turned on to activate Cry2. Time resolution, 1.0 frame per second. Scale bar = 10 μm.

**Movie S2. SMN$^{Tud}$ forms clusters in the optodroplet assay.**

NIH-3T3 cells expressing SMN$^{Tud}$. At time 0.0, blue light is turned on to activate Cry2. Time resolution, 1.0 frame per second. Scale bar = 10 μm.

**Table S1. Constructs used in this study**

| Construct | Uniprot ID | Insert Sequence | Backbone | Promoter |
|---|---|---|---|---|
| mCh-Cry2 | N/A | None | pHR_SFFV-mCherry-Cry2 | SFFV |
| $FUS^{IDR}$ | P35637 | aa 1-214 | pHR_SFFV-mCherry-Cry2 | SFFV |
| $hnRNPA1^{IDR}$ | P09651 | aa 186-320 | pHR_SFFV-mCherry-Cry2 | SFFV |
| $SMN^{IDR\text{-}N}$ | Q16637 | aa 1-91 | pHR_SFFV-mCherry-Cry2 | SFFV |
| $SMN^{Tud}$ | Q16637 | aa 92-145 | pHR_SFFV-mCherry-Cry2 | SFFV |
| $SMN^{IDR\text{-}C}$ | Q16637 | aa 146-294 | pHR_SFFV-mCherry-Cry2 | SFFV |
| *dm*$Tudor^{Tud}$ | P25823 | aa 2315-2515 | pHR_SFFV-mCherry-Cry2 | SFFV |
| Aub3 | O76922 | 3x(NPVIARGRGRGRK) | pLV-EGFP | eIF4a |
| Aub3(KG) | O76922 | 3x(NPVIAKGKGKGKK) | pLV-EGFP | eIF4a |
| $Spf30^{Tud}$ | O75940 | 72-132 | pHR_SFFV-mCherry-Cry2 | SFFV |
| $Tdrd1^{Tud}$#2 | O9BXT4 | 541-600 | pHR_SFFV-mCherry-Cry2 | SFFV |
| $Tdrd1^{Tud}$#3 | O9BXT4 | 762-821 | pHR_SFFV-mCherry-Cry2 | SFFV |
| $Tdrd1^{Tud}$#4 | O9BXT4 | 990-1048 | pHR_SFFV-mCherry-Cry2 | SFFV |
| $Tdrd3^{Tud}$ | Q9H7E2 | 555-615 | pHR_SFFV-mCherry-Cry2 | SFFV |
| $Tdrd4^{Tud}$#1 | Q9BXT8 | 726-784 | pHR_SFFV-mCherry-Cry2 | SFFV |
| $Tdrd4^{Tud}$#4 | Q9BXT8 | 1479-1539 | pHR_SFFV-mCherry-Cry2 | SFFV |
| $Tdrd6^{Tud}$#5 | O60522 | 1033-1088 | pHR_SFFV-mCherry-Cry2 | SFFV |
| $Tdrd6^{Tud}$#6 | O60522 | 1352-1411 | pHR_SFFV-mCherry-Cry2 | SFFV |
| $Tdrd8^{Tud}$ | Q9BXU1 | 78-137 | pHR_SFFV-mCherry-Cry2 | SFFV |
| $Tdrd9^{Tud}$ | Q8NDG6 | 944-1004 | pHR_SFFV-mCherry-Cry2 | SFFV |
| $Snd1^{Tud}$ | Q7KZF4 | 729-787 | pHR_SFFV-mCherry-Cry2 | SFFV |

**Table S2. Antibodies used in this study**

| Name | Epitope | Source | Application & Dilution |
|---|---|---|---|
| mCherry Antibody | mCherry | Invitrogen (#PA5-34974) | WB (1:2000), IF (1:500) |
| Coilin (H-300) | Coilin | Santa Cruz Biotechnology (#sc-32860) | IF (1:200) |
| SYM10 | sDMA | Millipore Sigma (#07-412) | WB (1:1000), IF (1:100) |
| SYM11 | sDMA | Millipore Sigma (#07-413) | WB (1:1000) |
| ASYM24 | aDMA | Millipore Sigma (#07-414 | WB (1:2000) |
| SMN (2B1) | SMN | Abcam (#ab5831) | IF (1:200) |
| Anti-green fluorescent protein, rabbit IgG fraction | GFP | Invitrogen (#A11122) | WB (1:2000) |
| GAPDH (FL-335) | GAPDH | SantaCruz Biotechnology (#sc-25778) | WB (1:2000) |
| Y12 | Sm | Gift from Joan Steitz | IF (1:10) |
| Anti-2,2,7-Trimethylguanosine Mouse mAb (K121) | TMG | Calbiochem (#NA02) | IF (1:200) |
| Rb pAb to SNRPC | U1 snRNP-C | Abcam (#ab82862) | IF (1:200) |
| CB7 | U1-70K | Gift from Doug Black | IF (1:200) |
| Anti-rabbit IgG Horseradish Peroxidase-Linked Species-Specific Whole Antibody | Rabbit IgG | GE HealthCare (#NA934) | WB (1:10,000) |
| Alexa Fluor 488-conjugated AffiniPure Donkey Anti-Rabbit IgG | Rabbit IgG | Jackson ImmunoResearch (#711-545-152) | IF (1:500) |
| Alexa Fluor 488-conjugated AffiniPure Donkey Anti-Mouse IgG | Mouse IgG | Jackson ImmunoResearch (#715-545-150) | IF (1:500) |

Abbreviations- WB: Western Blot, IF: Immunofluorescence
